# A Very virulent infectious bursal disease virus closely related to New York strain isolated from vaccinated small-scale broiler poultry farm in Addis Ababa, Ethiopia

**DOI:** 10.1101/2022.12.09.519830

**Authors:** Gizat Almaw, Abebe Olani, Melaku Sombo, Bekele Yalew

## Abstract

Infectious bursal disease virus (IBDV) was initially identified in the USA. The first IBD case in Ethiopia was reported in 2002 and since then vaccination was employed to control the disease. However, IBD outbreaks continued to occur despite the routine vaccination. The present study investigated IBD outbreak that had occurred in a vaccinated small-scale broiler poultry farm. Eight samples (four Bursa and four spleen) were collected and all were confirmed to be IBDV positive using reverse transcription-polymerase chain reaction (RT-PCR) targeting the hypervariable region of virus protein 2 (hVP2). All the IBDV isolates of this study were identified to be very virulent IBDV (vvIBDV) strains. For the four IBDV isolates the nucleotide and deduced amino acid sequence for hVP2 was determined. The nucleotide sequence identity of the VP2 gene of the present outbreak isolates (which showed 100% homology among themselves) with the previous 19 vvIBDV characterized isolates from Ethiopia ranged from 90.8% to 96.9% but greater than 96% identity was recorded with only six isolates. The deduced amino acid (aa) sequence of the present outbreak isolates contained aa residues commonly found in vvIBDV strains: A222, I242, I256, I294 and S299 suggesting the strains belong to genogroup three (G3). The phylogenetic analysis of the isolates showed that all isolates clustered separately from classical virulent, vaccine and variant strains and also distantly related to UK661 (UK) and DV86 (Netherlands) very virulent strains but unexpectedly closely related to the New York, USA strain. In conclusion, the present study reported vvIBDV strains (G3) isolated from a vaccinated broiler flock. Further research is needed to characterize the molecular epidemiology and pathogenicity of the vvIBDV strains and evaluate protective efficacy of the current IBD vaccine used for routine vaccination.

## Introduction

Infectious bursal disease (IBD) is an acute highly contagious disease of young chicken caused by infectious bursal disease virus (IBDV) that has lymphoid tissue as its primary target with special predilection for the bursa of Fabricius [1]. Since its original recognition in Gumboro, Delaware, and hence the synonym ‘Gumboro Disease’ [2] has resulted in economic losses in the poultry industry worldwide. IBD virus (IBDV) is a member of the genus *Avibirnavirus* of the family *Birnaviridae* and consists of two double-stranded RNA segments, designated as A and B [3, 4]. For IBDV strain classification, both phenotypic and genotypic methods were used and two serotypes designated as Serotype I and II [5] were identified. Serotype I virus was first described and is pathogenic for chickens. After classical IBD vaccines failed to elicit full protection and relatively high mortality and morbidity rates were observed in vaccinated flock, the emergence of a new variant strain of serotype I was investigated. This new variant was first described in Europe and designated as *very virulent strain-*vvIBDV [6]. The nomenclature for IBDV has changed over time due to the rapid genetic variation in hyper variable region of virus protein 2(hVP2). Based on pathogenicity and antigenicity, Serotype I was originally classified into four main pathotypes as: classical virulent (cv), antigenic variant (av), very virulent (vv), and attenuated (at) [7]. However, due to genetic variation a new nomenclature is proposed which classified IBDV into seven genogroups (G1-G7) [8]. According to this classification the traditional cv/atIBDV, avIBDV and vvIBV strains were classified as G1, G2 and G3, respectively and the remaining G4 to G7 are also described.

However, vvIBDV (G3) is still the most pathogenic strain and continues to be reported from different countries around the world including Ethiopia [9, 10, 11, 12, 13]. For Ethiopia, IBD is considered as an exotic disease, and is believed to be introduced with chickens imported from Europe. The first IBD case in Ethiopia was reported in 2002 after the outbreak that had occurred in broiler chickens at Debre Zeit [14]. By then this issue was a hot discussion topic in 2003 among participants of the 17^th^ Ethiopian Veterinary Association Annual Conference [15]. Now the IBD vaccine is produced in Ethiopia and is being used in routine vaccination programs. Despite routine vaccination, IBD outbreak continued to occur and is still a challenge for the poultry industry in Ethiopia. One possible reason for these outbreaks could be the IBDV genetic changes. These genetic changes can be assessed via detection and molecular characterization of mainly the hVP2 region of the IBDV genome, which is responsible for the virulence and antigenicity of IBDV [16]. To the best of authors’ knowledge other than the works of Negash et al., [13], Jenberie et al., [17] and Shegu et al. [18], similar studies of such kind were lacking in Ethiopia.

The purpose of this work was therefore to do molecular analysis and document isolates of IBDV obtained from field outbeak that had occurred in small-scale broiler poultry farm in Addis Ababa, Ethiopia.

## Materials and Methods

### Flock history

IBD outbreak had occurred in Addis Ababa, capital city of Ethiopia on June 9, 2015 in a small-scale broiler farm having a flock size of 2700 birds. The breed was Bovans white chickens, originated from one of the commercial poultry multiplication farms in Ethiopia distributing day-old-chicks across the whole of Ethiopia. At the time of the outbreak, the birds’ age was just two months. Within two days of the outbreak, 450 birds were died (16.7% mortality rate) and 650 birds became sick (24.1% morbidity rate). The birds were vaccinated for IBD via eye drop method at 1 - 2 weeks of age and the second vaccination was done at 3 - 4 weeks of age following the manufacturer instruction [19]. Each dose of vaccine contains live freeze-dried IBD (Intermediate Standard Strain) virus of 10^3^ EID50.

### Virus isolation by cell culture technique

Eight tissue samples (four bursa and four spleen) from four chicken were collected in virus transport medium (VTM) and submitted to Cell Culture Laboratory of the then National Animal Health Diagnostic and Investigation Center (NAHDIC), Ethiopia - now Animal Health Institute (AHI) and inoculated on to Vero cell line following the standard operating procedure of NAHDIC [20]. Briefly, tissue (sample) was put in a mortar and washed several times with phosphate-buffered saline (PBS) with antibiotics and anti-mycotic. Then using a coarse sand, the sample was triturated thoroughly and a 10% suspension made in PBS with antibiotic and antimycotic. The suspension was then centrifuged at 3000 rpm for 20 minutes and the supernatant was collected for inoculation. Monolayer Vero cell line with a confluence of > 70% was selected and the growth medium removed and washed twice with PBS. The specimen suspension was inoculated and incubated at 37°C for 60 minutes to allow the virus to adsorb to the cell culture. Then the maintenance medium (MEM with 2% FCS) was added and the flask was incubated at 37°C for seven days and the presence of the virus was detected by observing the cytopathic effect (CPE).

### RT-PCR

A real-time PCR was used to detect the presence of IBDV-specific VP2 genomic RNA in bursa’s and spleen samples of chickens. Bursa and spleen samples of each chicken were examined and these samples were put on Flinders Technology Associates (FTA) cards, and the FTA cards were sent to the GD Animal Health (Deventer, the Netherlands) for RNA extraction, reverse transcriptase (RT)-PCR and sequencing of the VP2 region.

### Method of RNA isolation from FTA cards

In total, 9 punches of the FTA cards were taken by using the FTA puncher, and these punches were used for the RNA extraction [27]. A volume of 270uL of RNA Rapid Extraction solution was added to the 9 punched FTA cards and shaken for 5 minutes. After shaking, a volume of 200μL was used for the RNA extraction using the MagMax method (Applied Biosystems). Positive and negative controls were included in each test run to control the quality of the PCR test run.

### RT-PCR procedure

For the RT-PCR, the following IBDV-VP2 primer sequences were used; 5’-GGT AGC CAC ATG TGA CAG-3’ (forward primer) and 5’-CGC TCG AAG TTR CTC ACC C-3’ (revere primer). For the amplification, the Light Cycler RNA Amplification SYBR Green-I kit (Roche) was used. Therefore, a volume of 3uL of RNA extract was mixed with 6.0 uL SYBR-Green (5x), 0.6uL RT-PCR enzyme mix, 3.6uL MgCl_2_, 0.5uL of the forward and the reverse primer (final concentration 0.33 μM) was mixed with 15.8uL of PCR grade water (RN-ase and RNA free water).

Samples were amplified using the following protocol; 30 min at 52°C followed by 30 seconds at 95°C (one cycle for cDNA production). The cDNA was amplified using 40 cycles of: 5 seconds 95°C, 10 seconds 57°C, and 30 seconds at 72°C. The presence of specific IBDV VP2 genomic sequences was examined using melting curve analyses.

### Sequence analyses

Positive RT-PCR products were collected for sequencing. Therefore, a volume of 15μL of PCR product was mixed with 15μL of PCR buffer (1x), and the same reverse and forward primers were used as described above for sequencing using the Sanger method.

### Phylogenetic analysis

Nucleotide sequence and the deduced amino acid sequence relatedness among the isolates themselves and with previous isolates was analyzed using BLAST in NCBI database search (https://blast.ncbi.nlm.nih.gov/Blast.cgi#) [28]. Phylogenetic tree was constructed using the sequenced hVP2 gene of the isolates together with 66 other sequences retrieved from GenBank (Supplementary Table S1) representing different origins and the seven genogroups of IBDV. Of these 66 isolates, 24 were isolates from Ethiopia and one vaccine strain produced and used locally (Ethiopia) for routine vaccination was also included in the analysis. The phylogenetic analysis was conducted using MEGA 11 software [21] using the maximum likelihood method, Tamura-Nei model (TN93) model, muscle alignment and 100 bootstraps. The bootstrap values are indicated at each node of the phylogenetic tree. The tree was edited using Fig Tree v1.4.4 software [29].

## Results

### Major clinical and postmortem findings

The major clinical findings observed in sick birds at the time of the outbreak were depression, severe fresh bloody diarrhea, shivering and rough hair coat. At postmortem examination, hemorrhages were seen on the thigh, breast muscle and under the skin (Fig.1). Bursa was swollen, hemorrhagic and edematous. Spleen was also enlarged. No histopathology was done.

**Fig. 1.**
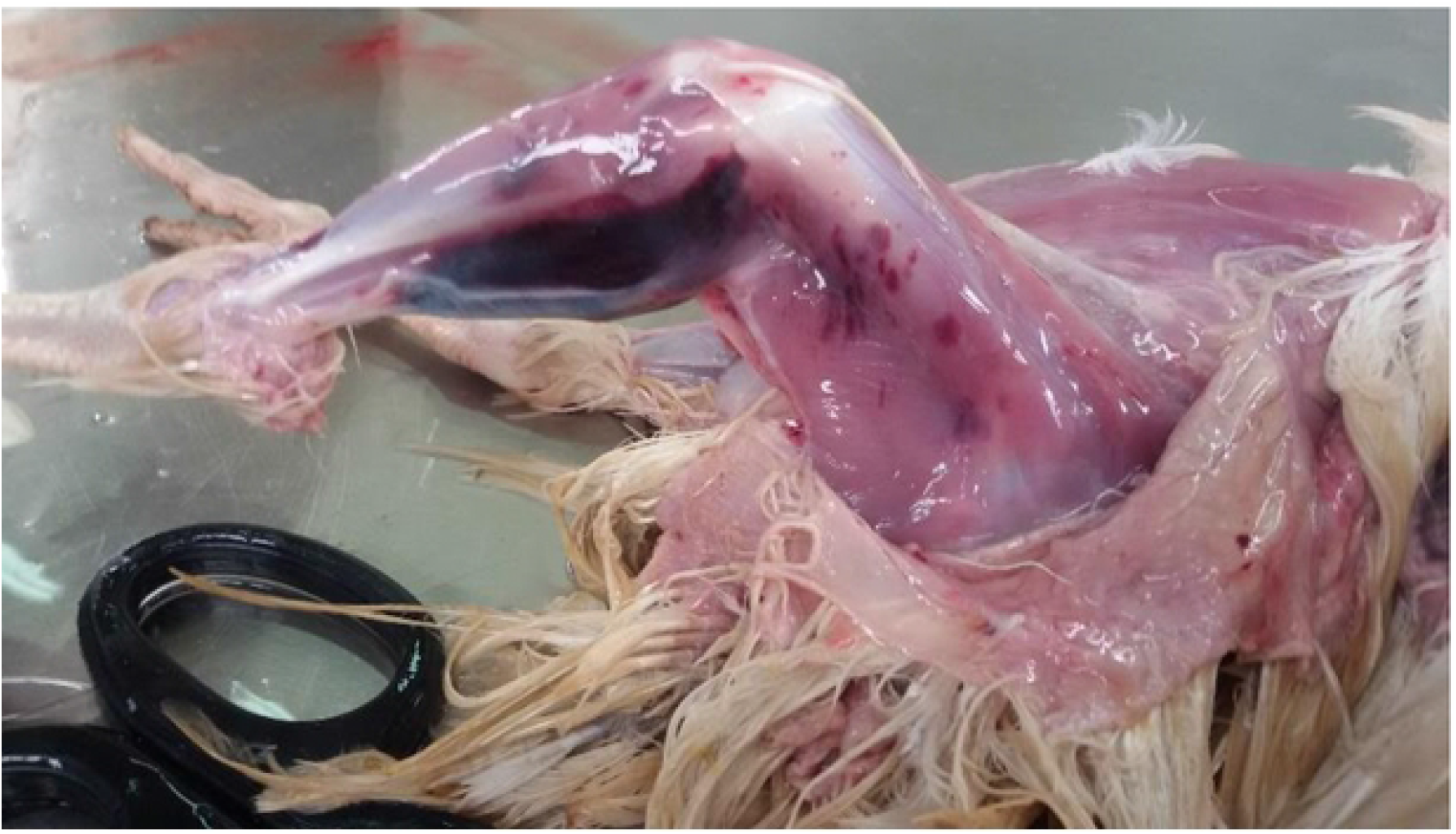
Infectious bursal disease. Hemorrhages on the thigh and under the skin.

### Virus isolation and RT-PCR test

All samples (n=8) showed cytopathic effect (CPE) at day five of the first passage. CPE was detected using inverted microscope by the presence of round and clumping cells where few were detached from the wall of the cell culture flask and float in the medium. Samples that were positive for CPE were run for RT-PCR targeting the VP2 gene of IBDV and all bursa and spleen samples were positive.

### VP2 sequencing

Among the RT-PCR positives, four (Busa samples) were sequenced and the nucleotide sequences of the four amplified products (423 nucleotides), showing 100% homology, were: 5’-GCAGCCGATGATTACCAATTCTCATCACAGTACCAAGCAGGTGGGGTGACGATCACACTGTTCTCAGCCAACATCGATGCCATCACAAGTCTCAGCATCGGGGGAGAACTTGTGTTCCAAACAAGCGTCCAAAGCCTCATACTAGGTGCTACCATCTACCTCATAGGCTTTGATGGGACTGCAGTAATCACCAGAGCTGTGGCCGCAGACAATGGGCTAACGGCCGGCACTGACAACCTTATGCCATTCAATATAGTGATTCCCACCAGCCAGATAACCCAGCCCATCACATCCATCAAACTGGAGATAGTGACCTCCAAAAGTGGTGGTCAGGCTGGGGATCAGATGTCATGGTCAGCAAGTGGGAGCCTAGCAGTGACGATCCACGGCGGCAACTACCCAGGGGCCCTCCGTCCCGTCACA.

The deduced amino acid sequences of the four sequenced samples (141 amino acids), showing 100% homology,were: AADDYQFSSQYQAGGVTITLFSANIDAITSLSIGGELVFQTSVQSLILGATIYLIGFDGTAVITRAVAADNGLTAGTDNLMPFNIVIPTSQITQPITSIKLEIVTSKSGGQAGDQMSWSASGSLAVTIHGGNYPGALRPVT.

The partial sequences of the VP2 gene of four strains detected in this outbreak were submitted to GenBank and accession numbers obtained. Accession numbers for isolates AA1/Eth/2015, AA2/Eth/2015, AA3/Eth/2015 and AA4/Eth/2015, respectively are OP727924, OP727925, OP727926 and OP727927.

Based on the obtained sequences, all the four samples were classified as very virulent (vv)-IBDV strains. The nucleotide and amino acid (aa) sequences were compared with the previously characterized IBDV strains (S1 Table) originated from Ethiopia, Africa, USA, Europe, Latin America and Asia. To this end the deduced amino acid sequence (aa 210-350) of VP2 of the present outbreak isolates was compared to vaccine strain (IBDV D78 and NVI Vaccine-Intermediate Standard Strain -Ethiopia), classical virulent IBDV strain (F52/70), and vvIBDV isolates that included: Netherlands DV86 (accession number: Z25482), UK661 (accession number: NC_004178), Russia 716 (accession number: MF142563), USA 1054NY/18 (accession number: MK408979), Washington St_G3b (accession number: MF142539) and Ethiopian vvIBDV isolates (Supplementary Table S2). All isolates of the present outbreak showed 90.7% and 91.5% aa identity to NVI Vaccine-Intermediate Standard Strain (Ethiopia) and strains of IBDV D78, respectively. All isolates had 100% aa similarity to all previous Ethiopian vvIBDV isolates selected for this comparison. The identity of classical virulent IBDV isolates (F52/70), vvIBDV isolates that includes Netherlands DV86, UK 661, Russia 716, USA 1054NY/18 and Washington St_G3b was 95.7%, 97.8%, 98.6%, 94.3%, 99.3%, and 97.9%, respectively.

The deduced amino acid sequence of the present outbreak isolates contained aa residues commonly found in vvIBDV strains: A222, I242, I256, I294 and S299. The characteristic motifs 222T, 249K, 286I, and 318D which are typical of the variant strain were not detected in any of the isolates. They differed from the prototype Netherland very virulent strain DV86 (accession number: Z25482) at three positions G254S, E300Q and A335V but they differed from the UK661 (accession number: NC_004178) type strain at two positions G254S and E300Q (Table 1). They differed only at a single position T270A with New York, USA Y14958 (accession number: MK408979) strain.

**Table 1.**
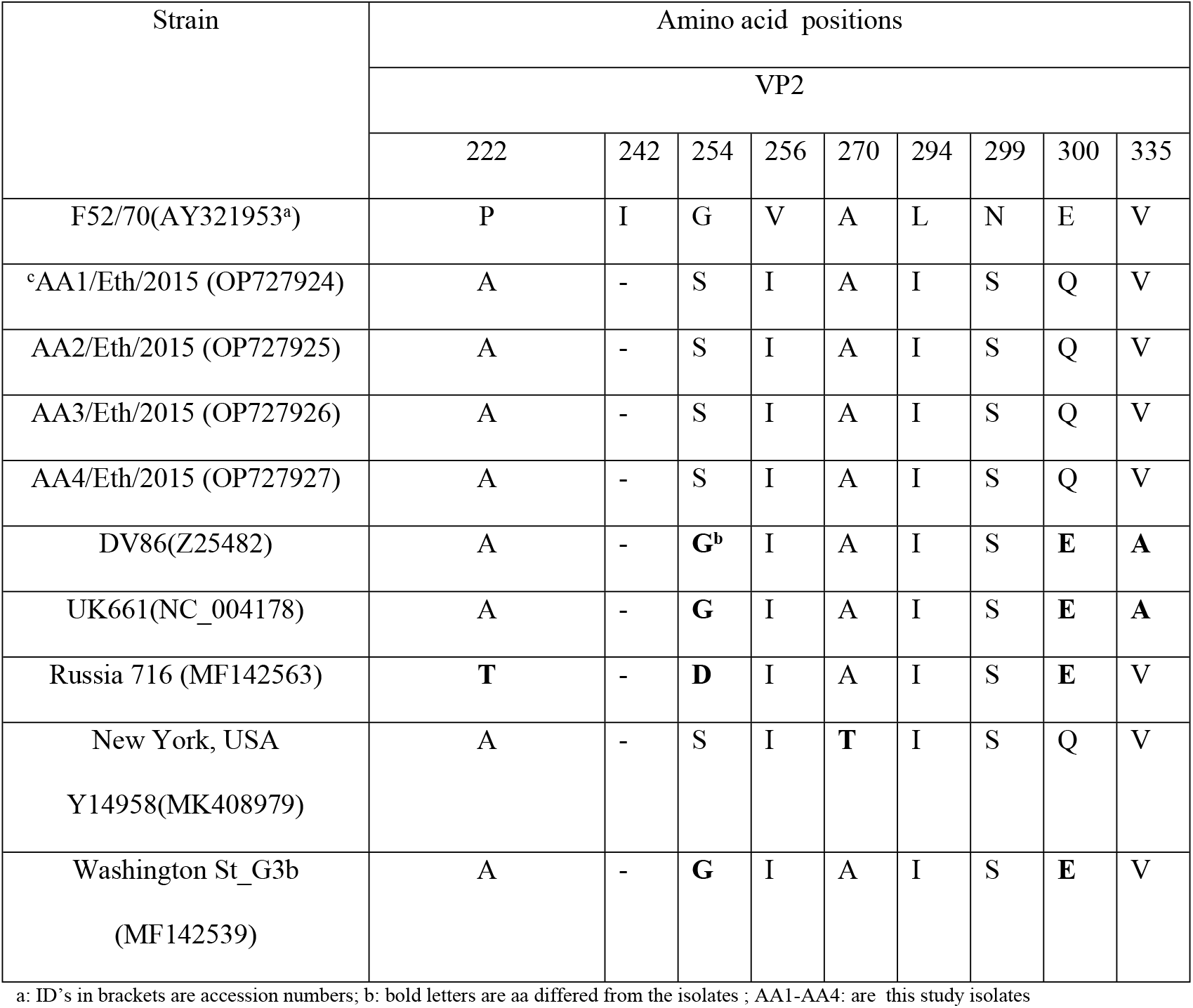
Comparison of key amino acid residues in the hypervariable region of VP2 of isolates, selected vvIBDVs and reference strains

| Strain | Amino acid positions |  |  |  |  |  |  |  |  |
| --- | --- | --- | --- | --- | --- | --- | --- | --- | --- |
|  | VP2 |  |  |  |  |  |  |  |  |
|  | 222 | 242 | 254 | 256 | 270 | 294 | 299 | 300 | 335 |
| F52/70(AY321953<sup>a</sup>) | P | I | G | V | A | L | N | E | V |
| °AA1/Eth/2015 (OP727924) | A | - | S | I | A | I | S | Q | V |
| AA2/Eth/2015 (OP727925) | A | - | S | I | A | I | S | Q | V |
| AA3/Eth/2015 (OP727926) | A | - | S | I | A | I | S | Q | V |
| AA4/Eth/2015 (OP727927) | A | - | S | I | A | I | S | Q | V |
| DV86(Z25482) | A | - | **G<sup>b</sup>** | I | A | I | S | **E** | **A** |
| UK661(NC_004178) | A | - | **G** | I | A | I | S | **E** | **A** |
| Russia 716 (MF142563) | **T** | - | **D** | I | A | I | S | **E** | V |
| New York, USA<br>Y14958(MK408979) | A | - | S | I | **T** | I | S | Q | V |
| Washington St_G3b<br>(MF142539) | A | - | **G** | I | A | I | S | **E** | V |

a: ID's in brackets are accession numbers; b: bold letters are aa differed from the isolates ; AA1-AA4: are this study isolates

The nucleotide sequence identity of the VP2 gene of the present outbreak isolates (which showed 100% homology among themselves) was compared to the VP2 sequence of previous 19 vvIBDV isolates from Ethiopia (GenBank) (S1 Table) collected at different time periods and areas. The similarity ranged from 90.8% to 96.9% but greater than 96% identity was seen with only six isolates. The nucleotide sequence similarity with the vaccine produced and used in routine vaccination (Intermediate Standard Strain) in Ethiopia (accession number: JQ684022) was 90.1%. The nucleotide sequence of the present outbreak isolates were also compared to the prototype very virulent strain DV86 of The Netherlands (accession number: Z25482) given the commercial poultry in central Ethiopia used to import day old chickens (DOC) and parent stock from The Netherlands and the similarity was 95.3%.

The phylogenetic analysis of all isolates of this study (red highlighted Taxa in in Fig. 2) and previously characterized Ethiopian vvIBDV isolates retrieved from GenBank (S1 Table) (light green colored Clades in Fig 2) clustered together though sublineages (four) present within each cluster indicating the existence of within diversity. Geographically the Ethiopian isolates including the present study isolates clustered together with the African isolates particularly with that of Nigerian and Kenyan isolates (blue colored Clades in Fig 2). Outside of Africa, the Ethiopian isolates revealed close relatedness to the U.S. 1/chicken/USA/1054NY/18 isolate (brown highlighted Taxa in Fig 2); however, not clustered with the vvIBDV strain of Europe such as UK661 and Netherlands DV86. By genogroup the present and previous Ethiopian isolates (except those clades colored yellow Fig 2) analyzed corresponds to the genogroup3 (G3) which is vvIBDV strain. The five previous Ethiopian isolates with clades colored yellow in Fig 2 clustered together with the classical attenuated vaccines strain D78 (highlighted in pink, Fig 2).

**Fig 2.**
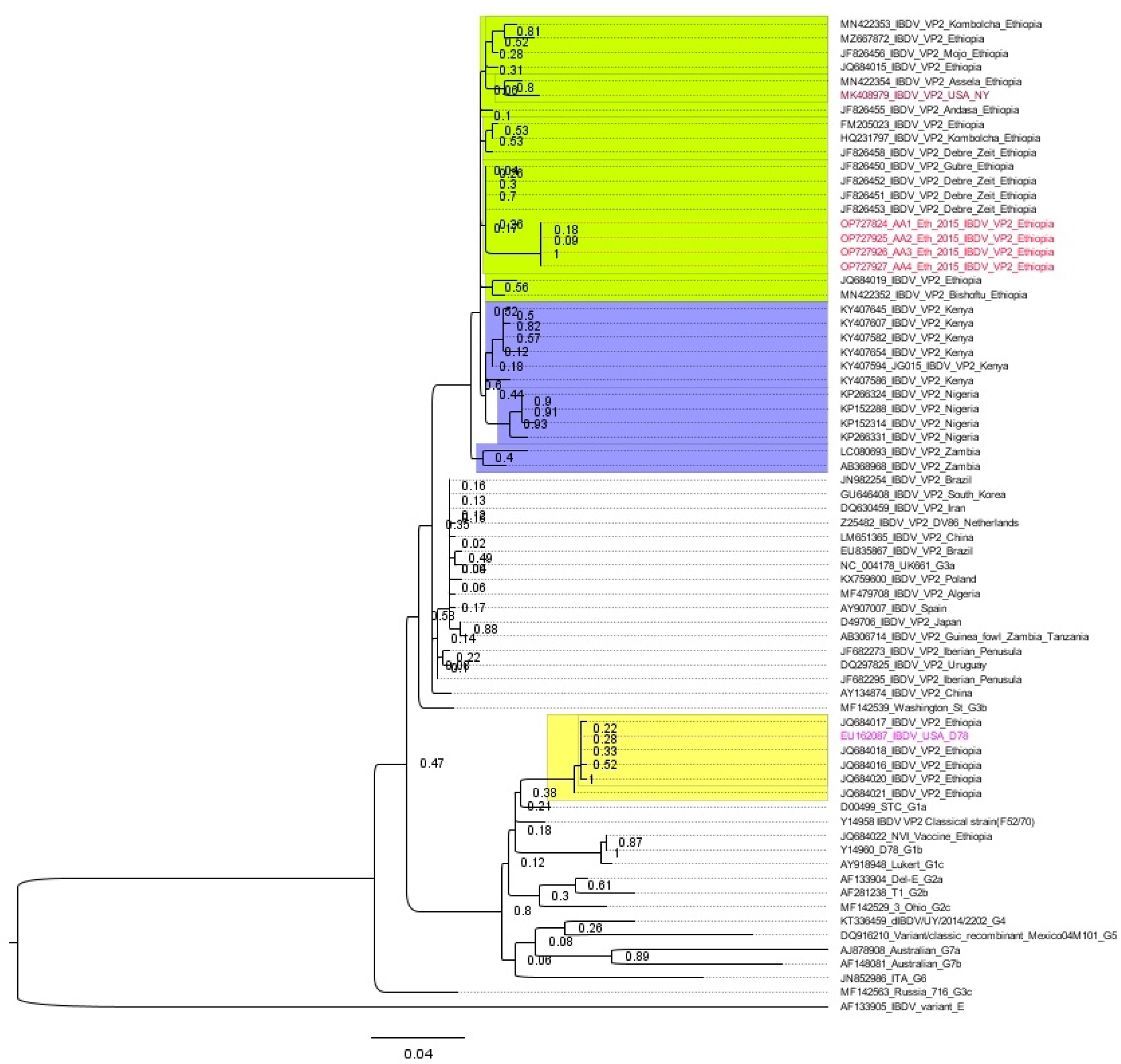
Phylogenetic tree produced using MEGA 11 software program [21] displaying the relationships of the present study isolates to vvIBDV isolates (U.S. 1/chicken/USA/1054NY/18, U.K 661, The Netherlands DV86, Ethiopian isolates retrieved from GenBank) and other genogroups designated as G1-G7 in the tree. In addition the phylogenetic tree described the relatedness of the isolates to African, European, Asian, Latin America/USA isolates. In the Taxa names all the first IDs followed by “_” sign represent the accession number for the respective isolates.

## Discussion

We did isolation and molecular characterization of IBDVs obtained from an outbreak that had occurred at small-scale broiler poultry farm in Addis Ababa, Ethiopia. The flock had a history of vaccination against IBD but experienced 24.1% morbidity and 16.7% mortality rate during the investigation period. Based on the RT-PCR test and obtained sequences, all isolates were classified as very virulent (vv)-IBDV strains. The deduced amino acid sequence contained the amino acid motifs: A222, I242, I256, I294 and S299 typical of vvIBDV strain suggesting the isolates belong to genogroup 3(G3) [23]. While vvIBDV strains first emerged in the late 1980s in Europe [24], the emergence of vvIBDV strains (vvIBDV) in Ethiopia is relatively a recent development. The first confirmed vvIBV strain was published in 2012 by Negash et al. [13] by analyzing isolates from the year 2005 to 2008.

A vvIBDV pathogenicity experiment study by Michel et al. [25] using isolate from a pullet flock in New York State showed 100% morbidity and 68.7% mortality within 4 days after a challenge to four-week old specific-pathogen-free (SPF) layer chicks. In Ethiopia Shegu and colleagues [18] recorded mortality rates as high as 80 %(n=4000) due to natural infection of vvIBV strain in an outbreak that had occurred in Sululta a place adjacent to the present outbreak. It would be interesting to undertake similar challenge studies using isolates from Ethiopia given Michel et al. [25]study indicated the isolate i.e. used for the experiment (accession no: MK408979) was most closely related to isolates from Ethiopia and suggested it is a new introduction into the U.S. We compared this isolate which is designated as New York 1/chicken/USA/1054NY/18 strain to the present outbreak isolates and it had 99.3% deduced aa and 95% nucleotide sequence identity in support of the previous observation. However, this needs epidemiological data to further analyze the possibility of transmission or introduction.

Nucleotide sequence and phylogenetic analysis of the hyper variable region of the VP2 gene showed that all isolates clustered separately from classical virulent, vaccine and variant strains and also distantly related to UK661 (UK) and DV86 (Netherlands) very virulent strains (Fig 2). And hence this study did not support Shegu et al. [18] study which suggests a possible introduction of the virus into Ethiopia from Europe, as poultry breeding stock were mainly imported from central European countries. All isolates of this study and 19 previously characterized Ethiopian vvIBDV isolates retrieved from GenBank and the U.S. 1/chicken/USA/1054NY/18 clustered together despite sublineages (four) present within each cluster indicating the existence of within diversity. This phylogenetic analysis also revealed isolates from Africa clustered together compared to Asian/Latin America/Europe though this requires large data to suggest any localized transmission. In this phylogenetic analysis we have tried to include all previous Ethiopian IBDV isolates available and they did not show any spatial clustering when we look at the origin of these isolates. Whether this absence of localization means the same genotype was circulating probably from a common source needs to be supported by epidemiological data including GPS data for mapping and to evaluate the routes of transmission. A study in Ethiopia by Negash et al. [13] described the dissemination of vvIBDV isolates of a clonal type in the chicken population in Ethiopia. They mentioned a probable reason for the rapid dissemination of IBDV may be the transportation of chickens from infected commercial farms to the breeding centers and from the breeding centers to farmers. This seems plausible reason even in the current poultry production system today where breeding centers are not required to certify their farm for any disease including IBDV despite they are sources for day old chicks (DOC) and pullets to farmers. Also Shegu et al. [18] and Jenberie et al. [17] recorded the clustering of Ethiopian vvIBDVs isolates where Ethiopian vvIBDVs showed considerable genetic homogeneity (0.0–2.6%), with phylogenetic analysis suggesting a single origin. However, Jenberie et al. [17] also reported vaccine strain unlike other previous studies in Ethiopia (13, 18) where six of the 10 Ethiopian IBDV isolates were phylogenetically related to classical attenuated vaccine IBDV D78, despite being detected from bursae with significant gross pathology of the bursa.

All previous confirmed vvIBDV isolates including the present study [13, 17, 18] were all obtained from outbreaks at different time points and places; however, only Negash et al. [13] included the vaccination history of the flocks where three of the 11 vvIBDV isolates were from a vaccinated farm. The present vvIBDV isolates were also from a vaccinated small scale broiler poultry farm. In Malaysia Aliyu et al. [22] isolated 11 IBDV strains where seven of them were vvIBDV from broiler flocks with a history of vaccination against IBD (mortality of <8%). There was no difference in aa residues between the present vvIBDV isolates and the vaccine strain (Intermediate Standard Strain) at the major hydrophilic peak B (aa residues from 314– 324) of the hVP2 and this vaccine is used for routine vaccination in Ethiopia. This is in agreement with Negash et al. [13] where they suggested the vaccination failure may not be due to changes in the viral antigenicity at this region of VP2. However, Ramon et al. [26] suggested possibility of the emergence of vaccine-escape variants due to replication of vvIBDV in vaccinated flocks in their multicentric study involving 481 farms in 11 European countries to evaluate the efficacy of different vaccines in preventing field strain bursa colonization. In this vaccine evaluation study, of all vaccinated farms (n=481), 10% of the samples tested positive for IBDV field strains, belonging to the genogroup three [26].

## Conclusions

In conclusion, the present study reported vvIBDV strains belonging to genogroup three isolated from a vaccinated broiler flock. The phylogentic analysis showed that the present isolates were clonal to the previous isolates in Ethiopia and New York, USA vvIBDV strain. This study also indicated vvIBDVs infection cannot be controlled using the current routine vaccination strategy and showed the need for further molecular analysis and pathogenicity studies.

## Data Availability Statement

The major findings of the study are presented in the article and additional information is included in the supplementary material. Accession numbers for isolates AA1/Eth/2015, AA2/Eth/2015, AA3/Eth/2015 and AA4/Eth/2015, respectively are OP727924, OP727925, OP727926 and OP727927.

## Author Contributions

Writing–original draft preparation was done by GA. Writing–review and editing was done by GA and AO. Phylogenetic analysis was done by GA. Investigation was done by GA, AO, MS, and BY. All authors have read and agreed to the published version of the manuscript.

## Ethics Statement

Ethical review and approval was not required as samples were collected as part of routine disease diagnosis/investigation following farmers’ request.

## Funding

Costs related to sample collection and consumables were covered by AHI as part of its annual budget dedicated for outbreak investigation.

## Conflict of Interest

The authors declare that there are no conflicts of interest.

## Acknowledgments

We thank Dr Dereje Kebede from Shola Veterinary Laboratory, Addis Abeba for facilitating the outbreak investigation at the farm. We are also grateful to the then National Animal Health Diagnostic and Investigation Center (NAHDIC), now renamed as Animal Health Institute (AHI), Sebeta, Ethiopia for the support of field logistics and supply of consumables. The authors also thank Mrs Menbere Kidane for her technical support in cell culture. We would like also to extend our appreciation to Dr Gerard Welberg, GD Animal Health, The Netherlands for doing the RT-PCR and sequencing.

## Supplementary Material

**S1 Table**: Accession numbers and Origins of IBDV sequences retrieved from GenBank and used for the phylogenetic analysis.

**S2 Table**: Identity of deduced amino acid sequences of the VP2 hypervariable region of IBDV strains detected in the study and reference and selected strains.

## Notes

### Competing Interest Statement

The authors have declared no competing interest.

